# Aggressive interference as a strategy for enhancing resource gain

**DOI:** 10.1101/2025.01.15.633229

**Authors:** Chet F. Rakocinski

## Abstract

Adaptive aggressive behavior should serve as a leverage mechanism for boosting resource gain above what would be obtained under the premise of equal access. The functional response for a consumer within a group of consumers should decline proportionately with the number of non-aggressive consumers vying for the same resources. The expression of aggressive interference should correspond with the potential for resource gain as determined by tradeoffs involving leverage, resource availability and consumer density. Models proposed in this paper expand on these assertions to make specific falsifiable predictions pertaining to the expression of aggressive interference with respect to the tradeoffs. Models predict the resource concentration at which aggressive interference should peak, as well as the functional response value and corresponding threshold resource concentration above which aggressive interference should cease. A mechanistic interpretation of the maximum proportion of time dedicated to aggressive interference can be expressed in terms of leverage and consumer density. Normalized equations for an aggressive agent along with any number of other consumers additively compose joint consumption under sustained restricted resource availability. Product log solutions of Rogers’ Random Predator Equation heuristically predict joint consumption by an aggressive agent along with any number of other consumers under resource depletion.

## Introduction

### Adaptive aggressive behavior

Adaptive explanations of biological traits refer to their functional design. Accordingly, adaptive behavior entails goal-directed orientation to environmental cues, as perceived through sensory and cognitive capabilities (Dill 1983; Hills 2006); and mediates interactions with the environment to promote Darwinian fitness (Hamilton 1964). Consequently, natural selection favors behavioral traits that improve individual reproductive success (Klopfer and Hailman 1972). Heritable genotypic variability modulates the expression of behavioral traits at the population level (Benus et al. 1991; Duckworth 2009). Phenotypic plasticity in behavioral traits also occurs in response to the local environment (Chevin and Lande 2015). Likewise, the capacity to learn can improve the efficacy of behavioral traits within a changing environment (Kieffer and Colgan 1992; Fawcett et al. 2013; Jackson et al. 2013; Jackson et al. 2014). Over deeper time scales, selection for secondary traits may reinforce adaptive behavior (Zuk et al. 2014). For example, secondary morphological traits like larger size, horns, tusks, etc, may evolve to enhance the efficacy of aggressive behavior and increase fitness (King 1973). Agonistic behavior encompasses a range of interactions between dominant and subordinate animals, ranging from attack to threaten, avoidance and retreat (Scott and Fredericson 1951). When used for gaining priority of access to limited resources, assertive aggressive behaviors include attacking, rushing, displaying, signaling, and other forms of intimidation. Such behaviors collectively fall under the rubric of competitive interference (Peckarsky 1991). Henceforth, the term aggressive interference refers to such asymmetric behavior.

Aggression pervades the animal kingdom (Holekamp and Strauss 2016). The Darwinian interpretation implies aggressive behaviors fulfill adaptive functions (Lorenz 1967; Kuse and De Fries 1976). As such, aggressive behavior may serve in the defense of self, mates, and offspring. Aggressive behavior can also impose preemptive access to resources (Giraldeau and Caraco 2000; Pellegrini 2008; Holdridge et al. 2016). Preemptive access may apply to food, mates, or space, either within an unfixed area of discovery, a fixed territory, or a refuge (Kilgour et al. 2018). The prevalence of intraspecific aggression over interspecific aggression (Lorenz 1967) implies that the degree of aggression corresponds with potential resource overlap (Moore 1978). The lack of preemptive resource use within any given species indicates the exception that proves the rule. Species that do not vie for access to limited resources must occur under ecologically unfavorable conditions or such a strategy. Under proper ecological conditions, some animals even cooperate to enhance resource gain (Nilsson et al. 2007; Mazzolini and Celani 2020), although competition prevails. Moreover, phenotypic plasticity in the expression of aggressive behavior occurs universally (Moody and Ruxton 1996; Sirot 2000; Gibert and DeLong 2015). Aggressive interference behavior thus provides a flexible strategy for garnering resources within the context of variable social and environmental conditions (Nilsson et al. 2004; Giraldeau and Dubois 2008).

The flexible expression of aggression affirms its adaptive value. Resource availability mediates whether aggressive interference behavior can act to enhance resource gain (Nilsson et al. 2004; Rands et al. 2006; Peiman and Robinson 2010; Fattorini et al. 2018). Consumer density also modulates the efficacy of aggressive interference by imposing immediate demands on resources. Indeed, the competitive relationship may shift from exploitative to interference as consumer density increases (Holdridge et al. 2016). Agonistic behavior preempts the time available for other gainful behaviors, such as searching, pursuing, and handling resources (Rakocinski et al. 1983). Thus, unlimited resource availability should preclude the expression of aggressive interference behavior in favor of time spent consuming resources (Moody and Ruxton 1996; Sirot 2000). Time dedicated to aggressive behavior poses a tradeoff with time spent and energy gained while consuming resources. A proper balance between tradeoffs should reflect the efficacy of aggressive interference in enhancing resource gain. Aggressive interference should become ineffective as a means of enhancing resource gain whenever the realized net gain becomes lower than potential net gain in the absence of the behavior.

Spatially aggregated distributions of consumers often occur in nature (Giraldeau and Caraco 2000). Groups form as a predation defense strategy, or in response to local concentrations of food, potential mates, or patchy habitats. Consumers may also aggregate as an outcome of differential speed of movement with respect to local resource depletion (Richards et al. 2000). Aggregated spatial distributions predispose consumers to vie for limited resources. Thus, contention for limited resources occurs commonly among consumers within groups (Sirot 2000; Giraldeau and Dubois 2008). Aggressive interference can impose access to limited resources within aggregations (Holdridge et al. 2016). Likewise, the expression of aggressive behavior hinges on dispersion of resources with respect to consumers (Scott and Fredericson 1951). Conversely, aggression mediates the spatial dispersion of consumers with respect to shared resources (Gübel et al. 2021). For example, aggressive interactions can produce uniform spacing of consumers with respect to locally shared resources. On the other hand, dominant consumers may prevent access to resources within well-defined territories (Verner 1977; Jensen et al. 2005). Effects of aggression on the spatial dispersion of consumers also extend to the population level (Duckworth and Badyaev 2007; Brownscombe et al. 2012). Uneven demographic effects of agonistic interactions include dominance hierarchies, growth depensation, differential mortality, debilitating effects of stress and injury, mating failure, prey switching, feeding niche expansion, etc. (Winberg and Sneddon 2022). On a suitably large spatial scale, aggressive interactions may facilitate the formation of ideal-free distributions marked by concurrence between the spatial dispersions of consumers and their resources (Farnsworth and Beecham 1997; Meer and Ens 1997; Krivan 2008).

### Optimality vs Satisficing

Optimality theory offers a conceptual framework for testing mechanistic hypotheses about how well an adaptive goal-directed behavior conforms to the maximization of an underlying cost/benefit function (Moors et al. 2017). Although animals seldom behave optimally (Fawcett et al. 2013), optimality criteria serve as standards of comparison for evaluating performance (Ward 1992). Alternatively, the satisficing criterion offers a more general standard predicated on the concept of sufficient utility rather than optimality (sensu Simon 1955; Goodrich et al. 2000). As such, any level of utility above the minimum cost/benefit constraint threshold defined by the satisficing criterion qualifies as sufficient for deciding to act. Although some critics allege that the satisficing criterion is trivial and unfalsifiable (Nonacs and Dill 1993), the same costs, benefits, and trade-offs can be used to describe both satisficing and optimality criteria (Ward 1992). The satisficing criterion complements the optimality criterion because the larger conceptual space of potential adaptive strategies contains the optimal strategy. Thus, the cost/benefit value corresponding with a satisficing criterion delineates a profitability limit with respect to whether a behavior qualifies as adaptive. In contrast, optimality epitomizes the maximum attainable utility of enacting a given behavior. When properly framed, the satisficing criterion offers a practical standard for evaluating the utility of a behavior. Natural selection should preclude the expression of behaviors that do not meet or exceed the satisficing criterion. Framed in this way, hypotheses predicated on satisficing criteria should be testable. Moreover, the use of a satisficing criterion as opposed to an optimality criterion facilitates the comparison of alternative behavioral strategies (Ward 1992). Additionally, joint use of a satisficing criterion in connection with an optimality criterion can serve to critically assess the adaptive value of a behavior. Herein, the premise of equal access offers a satisficing criterion for gauging the efficacy of aggressive behavior as a leverage mechanism for increasing resource gain.

### Standard Functional Response Models

The Holling functional response model (Holling 1959a; Holling 1959b) is a cornerstone of evolutionary ecology (DeAngelis et al. 1975) while also fulfilling a vital role within optimal foraging theory (Abrams 1982). Functional response models feature within diverse subfields, including population and community ecology, optimal foraging theory, trophic ecology, allometric scaling, and metabolic ecology (Abrams 1980, 1982, 1990, 2010, 2022; Weitz and Levin 2006; Basset and DeAngelis 2007; Brose et al. 2008; Brose 2010; Basset et al. 2012; Gibert and DeLong 2015; Holdridge et al. 2016). Functional response models specify consumption by one or multiple consumers and pertain to sustained or depleting resource levels (Rosenbaum and Rall 2018). Routinely used to characterize feeding rates, functional response models can describe the ‘consumption’ of any kind of renewable resource. Although static parameters drive the original deterministic Holling disc equations (Skalski and Gilliam 2001), later renditions incorporate flexible parameters and ecological covariates, like body size, consumer-dependent effects including behavioral interactions of consumers, as well as stochasticity. Despite their wide use, functional response models have not fully addressed how aggressive behavior may serve as leverage for enhancing resource consumption.

A general functional response model unifies all three standard types of functional response models along a continuum of model forms (Real 1977). The Type I functional response describes a linearly increasing approach to the asymptotic saturation level, and applies to consumers that can simultaneously search for and handle resources, like spiders tending a web, or consumers that do not spend time handling resources, such as filter feeders. Because these strategies would seldom pertain to animals using aggressive behavior to boost their rate of resource gain, the Type I functional response will not be considered further. The standard Type II functional response describes a decelerating asymptotic curve with respect to increasing resource concentration. The asymptotic upper limit reflects the capacity of the consumer to process the resource (Holling 1959). The Type III functional response also reaches an asymptotic upper limit with respect to increasing resource concentration. However, the Type III functional response forms a sigmoid curve which accelerates up to an inflection point, after which it decelerates and ultimately levels-off with increasing resource concentration. The sigmoid shape of the Type III curve reflects effects of learning or experience with respect to increasing resource concentration.

Standard functional response models specify instantaneous rates of consumption with respect to sustained levels of resource availability, as opposed to consumption integrated over time with respect to depleting concentrations of resource availability. At low levels of resource availability, the rate of encounter, or attack (a), drives the functional response, while the time taken to consume the resource, or handling time (h), constrains the functional response at elevated levels of resource availability (Rosenbaum and Rall 2018). Because search time (Ts), is inversely related to the encounter rate, the functional response approaches its asymptotic value, or saturates, as the search time between encounters approaches zero. For all three standard models, the functional response ranges from vanishingly low to asymptotically high consumption across a wide range in resource availability. Under unlimited resource availability, the functional design of the organism limits the maximum attainable rate of consumption.

### Consumer-Dependent Effects

Consumer-dependent effects involve behavioral interactions such as interference or facilitation among members of a group (Nilsson et al. 2004, 2007). Accounting for effects of consumer-dependence behavior often helps explain the nature of functional responses. Even in the absence of direct behavioral interactions, consumer dependence becomes inevitable with increasing consumer density (Ginzburg 2000). Likewise, consumer dependence may emerge as resources concentrate. Based on a meta-analysis, Skalski and Gilliam (2001) found that consumer-dependent effects influenced functional responses in most of the studies they considered (i.e., 18 out of 19). Likewise, Abrams and Ginzberg (2000) documented that most functional response studies conducted in the laboratory involved consumer-dependent effects. Examples of consumer-dependent effects include interference such as mutual distraction among consumers (Hassell 1978), or prey-dependent effects, like the inhibitory effects of consumers on prey (Free et al. 1977). Other consumer-dependent models treat the ratio of consumer density to resource concentration as the primary driver of the functional response (Hassell and Varley 1969; Abrams and Ginzburg 2000). Consumer-dependent effects offer increased realism (Skalski and Gilliam 2001), although consumer dependent models typically lack explicit mechanistic explanations (Nillson et al. 2007).

Interference behavior among consumers constitutes the most common kind of consumer-dependence effect studied within functional response models. Functional response models typically integrate interference as a symmetric “mass-action” effect with respect to the density of interacting consumers (Van Der Meer and Smallegange 2009). Symmetric models incorporate interference as an exponential effect with respect to the abundance of consumers (sensu Hassell and Varley 1969). Such models generally assume that time spent on interference is exclusive of time spent consuming resources (Beddington 1975; DeAngelis et al. 1975), although some models treat interference as nonexclusive of consumptive behaviors (Crowley and Martin 1989). When exclusive, interference acts as a consumer-dependent inhibitory effect on the collective functional response. Interference has also been modeled as independent of or dependent on resource concentration (Moody and Ruxton 1996). If dependent, interference should wane as the functional response approaches saturation in connection with increasing resource availability (Skalski and Gilliam 2001).

Incorporating consumer-dependent behavioral effects into functional response models should improve their explanatory capabilities (Nillson et al. 2007). Functional response models often fit the data better with the inclusion of consumer-dependence (Skalski and Gilliam 2001). For example, symmetric interference lowers the collective functional response through the mass action effect of reducing the time available for obtaining resources (Hassell, M. P. 1978). But asymmetric aggressive interference also exemplifies consumer-dependent interactions (Abrams and Ginzburg 2000). Functional response models based on asymmetric interference can distinguish different modes of aggressive behavior used for enhancing resource gain (Nillson et al. 2004). Furthermore, consumers facultatively express aggressive interference, depending on changing ecological trade offs (Stillman et al. 1997; Goss-Custard et al. 1998, as cited in Sirot 2000). Indeed, the Dove-Hawk game theory model confirms the evolutionary stability of a flexible aggressive interference strategy (sensu Smith and Price 1973; Cowden 2012; Křivan and Cressman 2017). Moreover, a Dove-Hawk game theory model predicts that the expression of aggressive interference should relate inversely to resource availability and increase with consumer density (Sirot 2000). This expectation contrasts some functional response studies that show a nonlinear hump-shaped relationship in aggression intensity with respect to food concentration (Moody and Ruxton 1996). Clearly, further development of mechanistic consumer-dependent functional response models will enhance ecological knowledge.

### The Leverage Effect of Aggressive Interference

Aggressive interference potentially raises the realized rate of resource gain (Goss-Custard et al. 1998, Sirot 2000; Nillson et al. 2004). Thus, interference-aggression functional response models should account for costs and benefits of directed aggression when used as leverage to boost resource gain (Meer and Ens 1997). The premise of equal access in principle can serve as a satisficing criterion for comparison with the realized cost/benefit value associated with using aggression as leverage. Furthermore, energy/time (E/T) maximization (Schoener 1971) provides a cost/benefit goal for comparison with the equal access criterion.

In the proposed models, the leverage coefficient (λ) specifies the degree to which aggressive interference boosts the rate of resource encounter for the aggressive agent. The magnitude of λ reflects its efficacy for enhancing gain. Thus, λ drives the resource encounter rate relative to a baseline value of 1.0, signifying the equal-access rate of encounter within a group of P consumers. Additionally, λ varies continuously in connection with the rate of resource encounter. Importantly, the elevated functional response cannot exceed the asymptotic upper limit denoted by T/h, as set by inherent consumer-design constraints.

An aggressive consumer should elevate its resource gain by suppressing competing consumers within its ‘area of discovery’ (sensu Hassell and Varley 1969; Sirot 2000), or the area under its control. The ‘area of discovery’ has been formally defined as, “the area it (i.e., an animal) visits per unit of searching time” (Hassell and Varley 1969). In this paper, the term, ambit, refers to the effective area or volume of discovery within which the aggressive consumer can elevate its rate of encounter with a resource. The ambit may be spatially fixed or dynamic with respect to the consumer. It also provides the basis for normalizing resource abundance to obtain resource concentration within the models.

In standard functional response models, total time (T) includes the time spent obtaining resources, and handling time (h) denotes time spent consuming one unit of resource (e.g., one prey item). With increasing resource availability, the functional response approaches the asymptotic value defined by T/h, while search time (Ts) approaches zero. In such models, searching for resources occurs exclusively of time spent consuming resources (Free et al. 1977). Likewise, time spent on aggressive interference precludes time spent consuming resources, including search, pursuit, attack, and handling behaviors. So, time spent on aggressive interference implies a tradeoff (Sirot 2000), potentially compensated for by a higher rate of consumption in fulfillment of an energy maximization strategy (E/T) (Schoener 1971). Although leverage may entail successive one-to-one agonistic interactions as in the Dove-Hawk game theory model, the operational effect of elevating resource gain do es not require individual contestants. Suppression of consumption by multiple consumers may occur concurrently. In this manner, the leverage coefficient (λ ) represents the effect of aggressive interference for elevating the rate of consumption.

The functional response models formulated herein predict how intensively aggression should be expressed when effectively used as leverage to enhance consumption. Because functional response models embody relevant parameters for specifying consumption relative to resource availability and consumer density, they offer a suitable framework for considering the implications of aggression used as leverage to elevate resource encounter rates. For example, the proposed models predict how the magnitude of aggression should vary with respect to resource availability and consumer density. Countervailing effects of time spent on aggression rather than consuming resources help define threshold conditions for when aggressive interference should be expressed.

## Results – Model Description

### Type II functional response – leverage within a group under sustained resource availability

The standard Type II functional response hinges on a balance of the encounter rate and handling time relative to resource concentration (N). The difference equation defined for a single consumer recursively describes the Type II functional response over a fixed period:

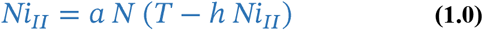

where,

Ni_II_ = number of resource units consumed;

a = rate of encounter per unit of resource per unit time (e.g., mates, food items, etc.);

N = abundance or concentration of resources;

T = total number of time units while obtaining resources (e.g., hrs);

h = handing time per unit of resource consumed.

When solved, equation **1.0** yields the standard Type II functional response in the form of the disc equation for a single consumer :

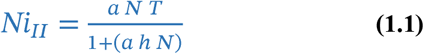

The disc form of the Type II equation describes a decelerating rectangular hyperbola with respect to increasing sustained resource concentration (N). The functional response approaches the asymptotic saturated level corresponding with the maximum of T/h as resource availability increases. The consumer has reached its full physical capacity at T/h because all available time is committed to handling resources. The model assumes a constant encounter rate, a, with respect to resource abundance. The individual consumes NiII resource units during the period, T, dedicated to obtaining resources at a sustained level of resource availability, N (i.e., N not depleted). This implies that resources are renewed by some process at the same rate that they are consumed. At low levels of N, the encounter rate governs the functional response, (i.e., the reciprocal of search time per unit of resource), and at high levels of N, handling time governs the rate of consumption. Thus, total search time, Ts = [T – (h × NiII)], and search time per unit of resource consumed, Ts/NiII = [T – (h × NiII)]/ NiII.

When multiple consumers co-occur within an unrestricted resource availability scenario, the foregoing Type II functional response for an individual will scale up directly with the abundance (or density) of non-overlapping consumers (P). The difference equation for multiple consumers facing unrestricted resources recursively describes the Type II functional response over a fixed period:

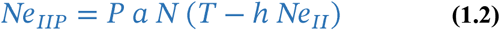

When solved, equation **1.2** yields the standard Type II functional response in the form of the disc equation for multiple consumers:

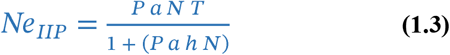

Thus, Ne_IIP_ stands for consumption by P consumers within an *unrestricted* resource availability scenario.

This multiple consumer-extended form of the Type II functional response implies each consumer reaches the same average rate of resource consumption defined by equation 1.1 in an additive manner (Fig. 1). Per capita consumption is independent of the presence of other consumers and resource availability stays at the level supporting consumption by P consumers collectively. Therefore, the encounter rate, a, stays undiminished within the ambit of each consumer. For example, a consumer-extended functional response might reflect consumption by multiple consumers across an expansive prey field. Under such an unrestricted resource availability scenario, aggressive interference would not effectively increase resource encounter rates.

**Figure 1.**
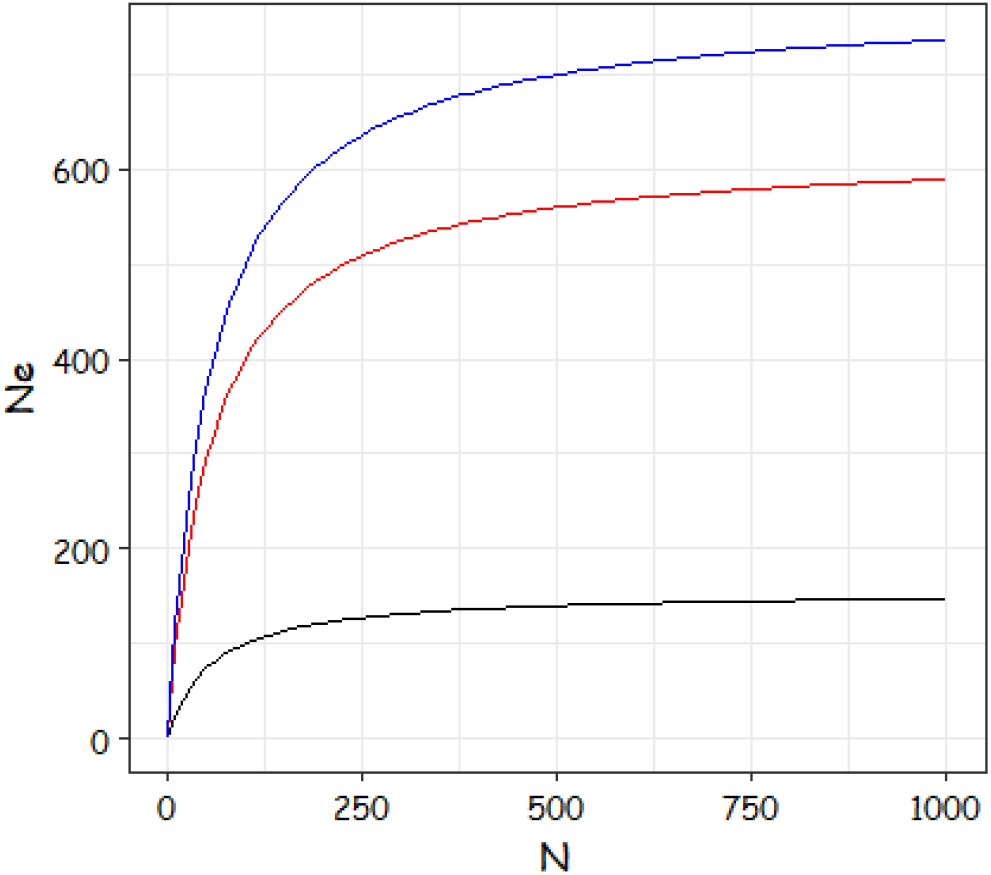
Type II functional responses for multiple consumers cooccurring within an *unrestricted* resource availability scenario. Functional responses scale up additively with the abundance of non-overlapping consumers. Black curve – P = 1, asymptotic Ne = 150; Red curve – P = 4, asymptotic Ne = 600; Blue curve – P = 5, asymptotic Ne = 750 (i.e., sum of black and red curves), a = 0.2, h = 0.09, T = 14 (P = number of consumers; Ne = consumption by P consumers; N = resource concentration; a = resource encounter rate; h = handling time per unit resource; T = total time).

This multiple consumer-extended form of the Type II functional response implies each consumer reaches the same average rate of resource consumption defined by equation **1.1** in an additive manner (Fig. 1). Per capita consumption is independent of the presence of other consumers and resource availability stays at the level supporting consumption by P consumers collectively. Therefore, the encounter rate, a, also stays undiminished within the ambit of each consumer. For example, a consumer-extended functional response might reflect consumption by multiple consumers across an expansive prey field. Under such an *unrestricted* resource availability scenario, aggressive interference would not effectively increase resource encounter rates.

When multiple consumers co-occur within a restricted resource availability scenario, the per capita encounter rate reflects overlapping resource use. This restricted model applies to a group of P consumers virtually sharing the ambit area/volume of a single consumer under an unrestricted resource availability scenario. Accordingly, multiple consumers partition the rate of resource consumption. This reduces the per capita rate of encounter for each consumer, while a resource replacement process sustains the restricted Type II functional response. The difference equation for a single consumer recursively describes the Type II functional response based on the premise of equal access:

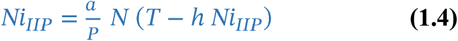

where,

a/P denotes the per capita encounter rate; and

Ni_IIP_ = per capita resource units consumed.

As a reflection of the virtual overlap, P may be an integer or a fractional number.

When solved, equation 1.4 yields the Type II functional response for an individual in the context of P consumers under restricted resource availability (Fig. 2):

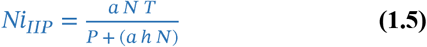

**Figure 2.**
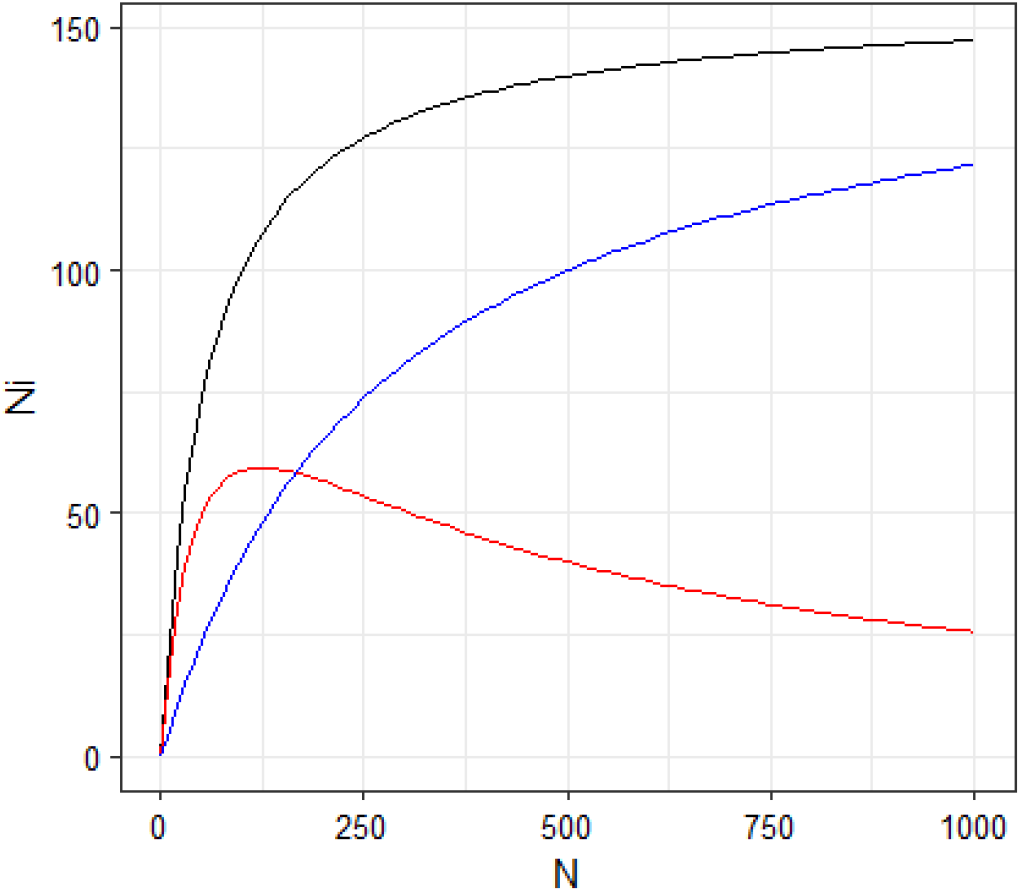
Type II functional responses for consumers within a restricted resource availability scenario. The difference in functional responses for a consumer with exclusive access to a resource (i.e., a = 0.2, h = 0.09, T = 14, black curve) and that for a consumer with equal access to the resource within a group (i.e., encounter rate = a/P; blue curve), impels the extent to which aggression can act as leverage for boosting consumption (i.e., red difference curve). Blue curve, P = 5, a/P = 0.2/5 = 0.04, h = 0.09, T = 14. Difference curve converges to zero at Ni_IIP_ = T/h for the blue curve (i.e., asymptote = 155.6). The peak value of the difference curve (i.e., maximum aggression intensity) occurs at N = sqr(P)/ah, (i.e., 124.23) (P = number of consumers; Ni = consumption by an individual consumer; N = resource concentration; a = resource encounter rate; h = handling time per unit resource; T = total time).

When multiple consumers co-occur within a restricted resource availability scenario, the Type II functional response will scale up with the abundance (or density) of overlapping consumers (P). The difference equation for multiple consumers facing restricted resources recursively describes the Type II functional response over a fixed period:

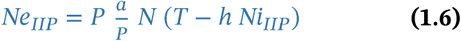

When solved, equation 1.6 yields the Type II functional response for multiple overlapping consumers under restricted resource availability:

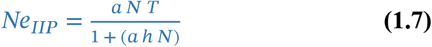

Where Ne_IIP_ stands for consumption by P consumers within a restricted resource availability scenario. The functional response equation for P consumers under restricted resource availability is identical to the standard per capita functional response equation described by equation 1.1.

Under restricted resource availability, a consumer may use aggressive interference to elevate its rate of encounter to any value above a/P, but not above a. Leverage (λ) signifies the effect of aggressive interference by a dominant consumer to elevate resource gain above the level inferred by the premise of equal access within a group of P consumers. Consumption cannot exceed the effect of P consumers, as λ/P = 1 implies the per capita maximum rate of consumption under unrestricted resource availability.

To sum up, The following conditions apply to the realization of λ:

1. λ must exceed 1 for aggressive interference to be gainful,
2. λ cannot exceed P (i.e., λ ≤ P),
3. λ varies continuously in connection with virtual overlap within the ambit.

Within the functional response of an aggressive consumer, the time spent on interference (i.e., T_λ_) as opposed to obtaining resources denotes a key tradeoff (c.f., Sirot 2000). The ensuing model assumes that time spent on aggressive interference is exclusive of time spent searching, pursuing, or apprehending/handling resources. Thus, the total time spent obtaining resources (i.e., T) comprises the time spent searching for and consuming resources in addition to that dedicated to aggressive interference (i.e., T_λ_). So, search time (i.e., T_s_) and handling time (i.e., T_h_) constitute Tf (i.e., T_f_ = T_s_ + T_h_), or T T_λ_. In this context, T_f_ represents time dedicated to obtaining resources at the leverage-enhanced rate of encounter for an aggressive consumer.

The following difference equation recursively defines the Type II functional response for an aggressive consumer within a group of consumers with equal access to resources:

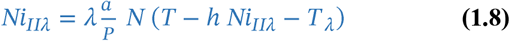

When solved, equation **1.8** yields the disc equation form of the Type II functional response for a consumer using aggressive interference as leverage :

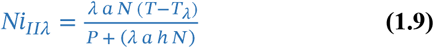

Ni_IIλ_ in equation **1.9** specifies the Type II functional response for an aggressive consumer overlapping with an effective group of P consumers at a sustained level of resource availability, N. The extent to which the functional response specified by equation **1.9** exceeds that for equation **1.5** at any level of resource availability (N), reflects the efficacy of aggressive interference behavior for elevating resource gain.

The difference between Ni_II_ (equation **1.1**) and Ni_IIP_ (equation **1.5**) defines how aggressive interference should perforce apply with respect to N. This follows the premise that the scope for elevating resource gain drives the degree of aggressive interference in terms of T_λ_. Thus, T_λ_ should reflect the normalized difference between Ni_II_ and Ni_IIP_ :

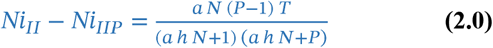

This difference describes a right-skewed hump-shaped curve with respect to N (Fig . 2). The proportional difference between Ni_II_ and Ni_IIP_ ranges between 0 at the lower and upper extremes and 1 at some intermediate level of N, for which the difference is greatest. The greatest difference occurs at Ni_IIm ax_, where the partial derivative of the difference curve with respect to N = 0 :

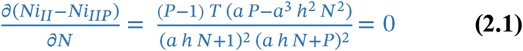

Assuming all parameters are positive:

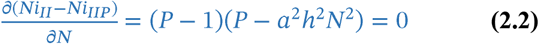

Solving Equation **2.2**:

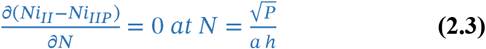

So: 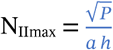, such that:

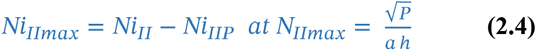

After substitution and rearrangement:

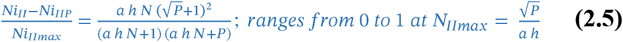

### Type II functional response – Threshold for aggressive interference

Based on the foregoing, the threshold resource concentration and corresponding functional response value at which aggressive interference should cease can be deduced. The threshold resource concentration for the expression of aggressive interference should concur with the intersection of the respective functional responses ( Fig. 3 ). Also, the intersection of Ni_IIP_ and Ni_IIλ_ corresponds with the threshold resource concentration, N_II crit_:

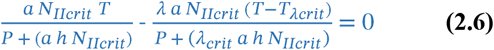

**Figure 3.**
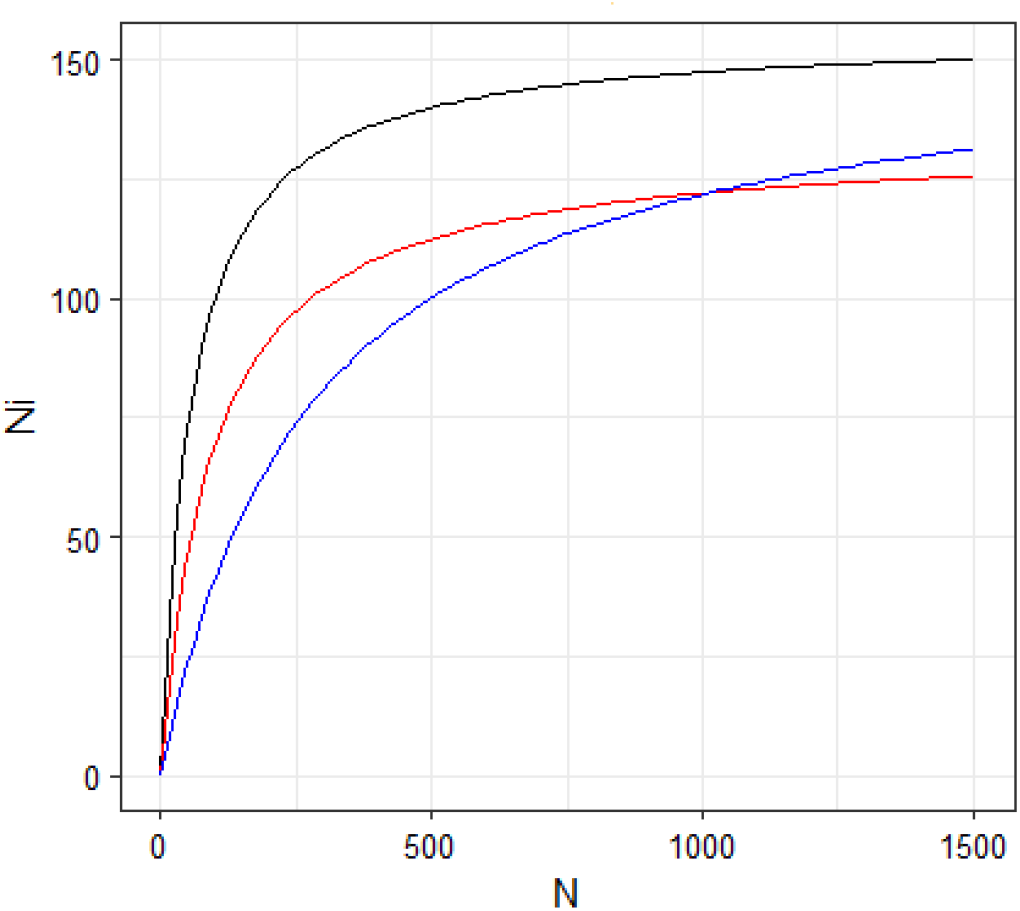
Type II functional responses for consumers within a restricted resource availability scenario, including that for an aggressive consumer exerting leverage on resource gain. Parameters in common: a = 0.2, h = 0.09, T = 14. The black curve describes the functional response of a consumer with exclusive access to the resource, while the blue curve describes the per capita functional response for one of five consumers (i.e., P = 5) with equal access to the resource (i.e., encounter rate = a/P). The red curve describes the functional response for an aggressive consumer applying leverage (i.e., λ = 3, T_λ_ = 2). Aggressive interference is expected to occur between resource levels N = 0 and N_IIcrit_ = 1,018.52, where the curves intersect at Ni_IIcrit_ = 122.22. (P = number of consumers; Ni = consumption by an individual consumer; N = resource concentration; a = resource encounter rate; h = handling time per unit resource; T = total time).

Solving equation **2.6** for N_II crit_:

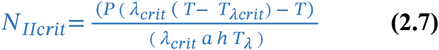

In equation **2.7**, the threshold time dedicated to aggressive interference (i.e., T_λcrit_) and threshold leverage (i.e., λ_crit_) are explicit . Solving equation **2.6** for these critical parameters yields the thresholds for time dedicated to aggressive interference (T_λcrit_ ) and leverage (λ_crit_):

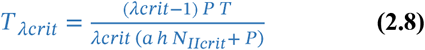

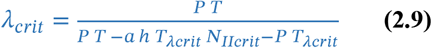

More generally, the product of T_λmax_ and the normalized difference between per capita functional response s for *unrestricted* exclusive access and *restricted* equal access consumers specifies T_λ_. Accordingly, for the Type II functional response:

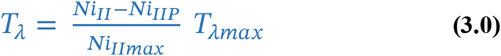

Upon rearrangement of equations 1.9 and 2.5:

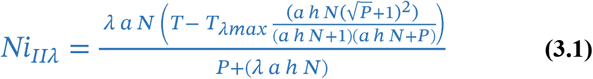

The threshold Type II functional response value (Ni_IIcrit_ ) corresponding to the threshold resource concentration (i.e., N_II crit_) for the expression of aggressive interference occurs where there is no difference between equations **3.1** and **1.5**:

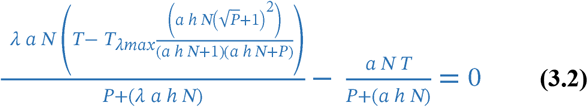

Solving expression **3.2** for Ni_IIcrit_ :

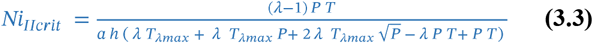

where Ni_IIcrit_ stands for the threshold functional response value for which no difference exists between aggressive interference and equal access strategies. Equation **3.3** obviates resource concentration. Thus, Ni_IIcrit_ denotes the point where aggressive interference and equal access functional responses intersect with respect to the threshold resource concentration, N_IIt_ (Fig. 3). Implicitly, profitability for the equal access strategy exceeds that for the aggressive interference strategy at resource concentrations greater than N_II crit_.

### Type III functional response – leverage within a group under sustained resource availability

The Type III functional response forms a sigmoid curve for which the rate of encounter systematically increases as an effect of experience. The difference equation defined for a single consumer recursively describes the Type III functional response over a fixed period:

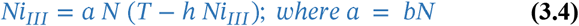

and thus,

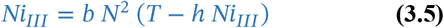

where,

Ni_III_ = number of resource units consumed;

b = the resource-concentration dependent rate of encounter (i.e., the attack rate, a = bN);

N = abundance or concentration of resources;

T = total time while obtaining resources (e.g., hrs);

h = handing time per unit of resource consumed.

When solved equation **3.5** yields the standard Type III functional response for a single consumer in the form of the disc equation:

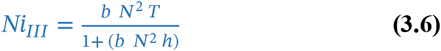

When multiple consumers co-occur within an unrestricted resource availability scenario, the foregoing Type III functional response for an individual will scale up directly with the abundance (or density) of non-overlapping consumers (P). The difference equation for multiple consumers facing unrestricted resources recursively describes the Type III functional response over a fixed period:

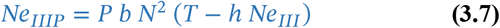

Thus, Ne_III_ stands for consumption by P consumers within an *unrestricted* resource availability scenario.

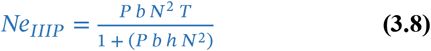

This multiple consumer-extended form of the Type III functional response implies each consumer reaches the same average rate of resource consumption defined by equation **3.6** in an additive manner (Fig. 4). Per capita consumption is independent of the presence of other consumers and resource availability stays at the level supporting consumption by P consumers collectively. Therefore, the encounter rate, b, stays undiminished within the ambit of each consumer. In nature, such a consumer-extended functional response might reflect consumption by multiple consumers across an expansive prey field. Under such an *unrestricted* resource availability scenario, aggressive interference would not effectively increase resource encounter rates.

**Figure 4.**
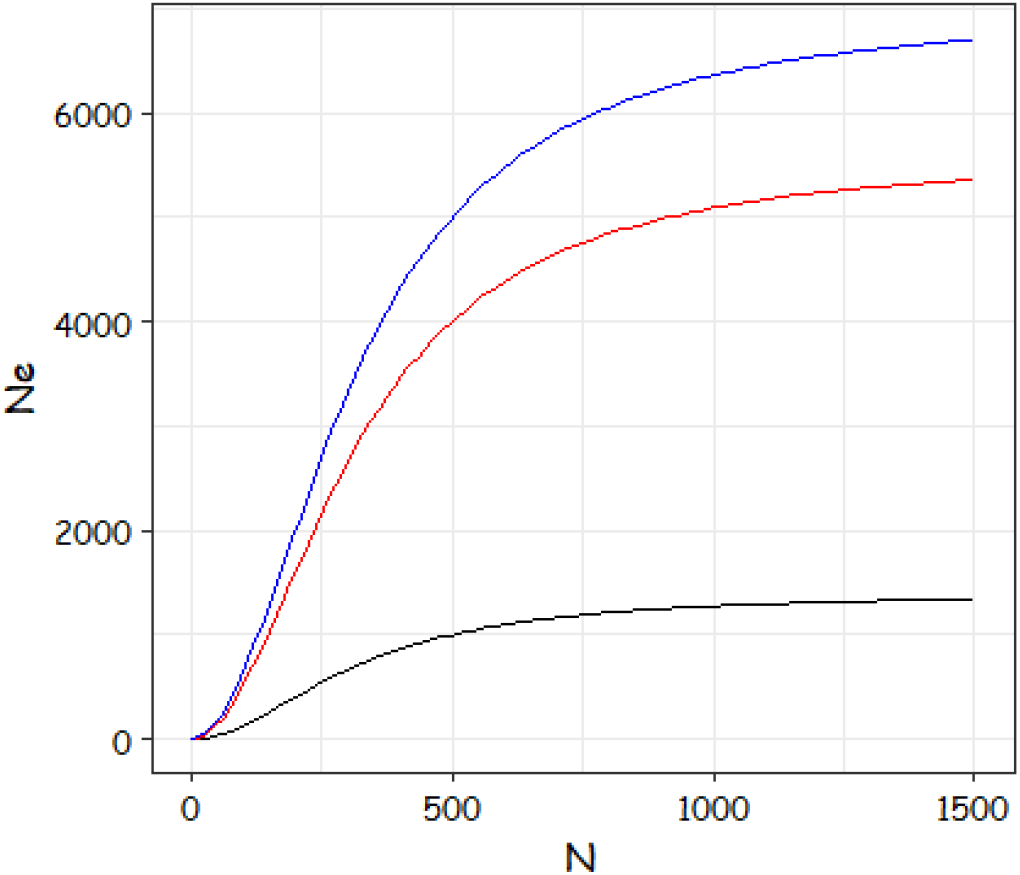
Type III functional responses for multiple consumers cooccurring within an unrestricted resource availability scenario. Functional responses additively scale up with the abundance of non-overlapping consumers (P). Black curve – P = 1, asymptotic Ne = 1400; Red curve – P = 4, asymptotic Ne = 5600; Blue curve – P = 5, asymptotic Ne = 7000 (i.e., sum of black and red curves), b = 0.001, h = 0.01, T = 14 (P = number of consumers; Ne = consumption by P consumers; N = resource concentration; b = resource encounter rate; h = handling time per unit resource; T = total time).

When multiple consumers co-occur within a *restricted* resource availability scenario, the per capita encounter rate reflects overlapping resource use. Thus, this model focuses on a group of P consumers virtually sharing the ambit area/volume of a single consumer under an *unrestricted* resource availability scenario. Accordingly, multiple consumers partition the rate of resource consumption. This reduces the per capita encounter rate for each consumer, while a resource replacement process sustains the *restricted* Type III functional response. The difference equation for a single consumer recursively describes the Type III functional response based on the premise of equal access:

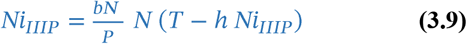

where,

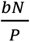 denotes the per capita encounter rate; and

Ni_IIIP_ = per capita resource units consumed.

As a reflection of the average overlap, P may be an integer or a fractional number.

When solved, equation **3.9** yields the Type III functional response for an individual in the context of P consumers under *restricted* resource availability (Fig. 5):

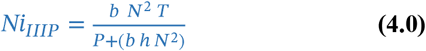

**Figure 5.**
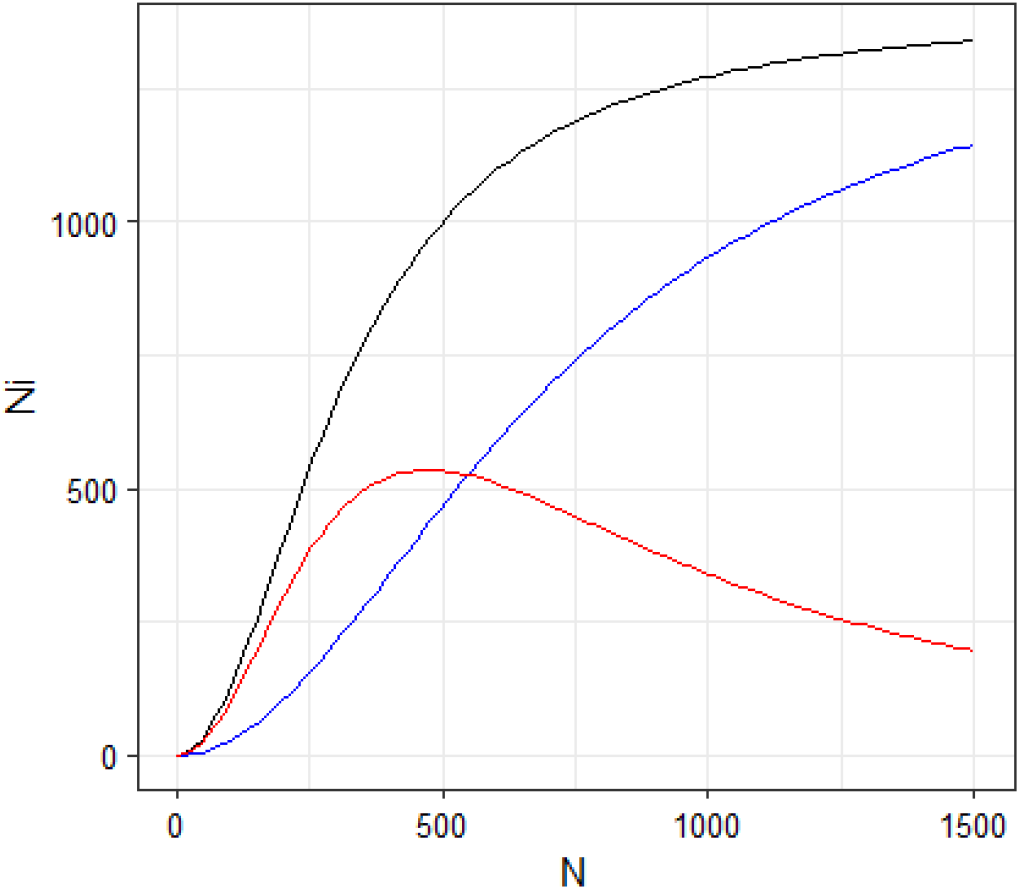
Type III functional responses for consumers within a *restricted* resource availability scenario. The difference in functional responses for a consumer with exclusive access to a resource (i.e., b = 0.001, h = 0.01, T = 14, black curve) and that for a consumer with equal access to the resource within a group (i.e., attack rate = (bN/P); blue curve), impels the extent to which aggression can act as leverage for boosting consumption (i.e., red difference curve). Blue curve, P = 5, bN/P = (0.001 N)/5; h = 0.01, T = 14. Difference curve converges to zero at N_iIIIP_ = T/h for blue curve (i.e., asymptote = 1400). The peak value of the difference curve (i.e., maximum aggression intensity) occurs at N = (frth(P)/sqr(b) sqr(h)), (i.e., 472.87). (P = number of consumers; Ni = consumption by an individual consumer; N = resource concentration; b = resource encounter rate; h = handling time per unit resource; T = total time).

When multiple consumers co-occur within a *restricted* resource availability scenario, the Type III functional response will scale up with the abundance (or density) of overlapping consumers (P). The difference equation for multiple consumers facing *restricted* resources recursively describes the Type III functional response over a fixed period:

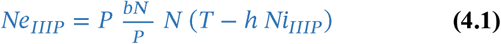

When solved, equation **4.1** yields the Type III functional response for multiple overlapping consumers under *restricted* resource availability :

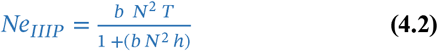

Where Ne_IIIP_ stands for consumption by P consumers. The functional response equation for P consumers under *restricted* resource availability is identical to the standard per capita functional response equation described by equation **3.6**.

The following difference equation recursively defines the Type III functional response for an aggressive consumer within a group of consumers with equal access to resources:

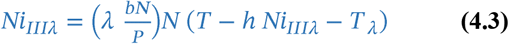

When solved, equation **4.3** yields the disc equation form of the Type III functional response for a consumer using aggressive interference as leverage:

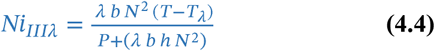

Ni_IIIλ_ in equation **4.4** specifies the Type III functional response for an aggressive consumer overlapping with an effective group of P consumers at a sustained level of resource availability, N. The extent to which the functional response specified by equation **4.4** exceeds that for equation **4.0** at any level of resource availability (N), reflects the efficacy of aggressive interference behavior for elevating resource gain.

The difference between Ni_III_ (equation **3.6**) and Ni_IIIP_ (equation **4.0**) defines how aggressive interference should perforce apply with respect to N. This follows from the premise that the scope for elevating resource gain drives the degree of aggressive interference in terms of T_λ_. Thus, T_λ_ should reflect the normalized difference between Ni_III_ and Ni_IIIP_:

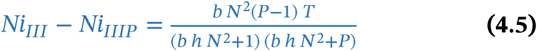

Like for the Type II model, this difference describes a hump shaped curve with respect to N (Fig. 5). The proportional difference between Ni_III_ and Ni_IIIP_ ranges between 0 at the lower and upper and 1 at some intermediate level of N, for which the difference is greatest. The greatest difference occurs at Ni_IIImax_, where the partial derivative of the difference curve with respect to N = 0:

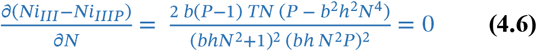

Solving equation **4.6**:

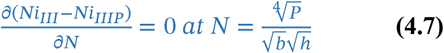

So 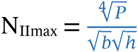, such that:

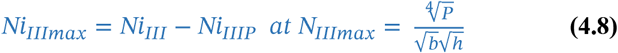

After substitution and rearrangement:

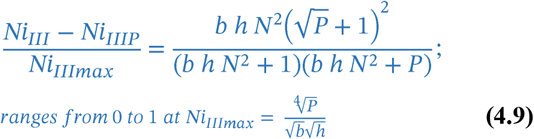

### Type III functional response – Threshold for aggressive interference

As for the Type II functional response, the threshold resource concentration and corresponding Type III functional response value at which aggressive interference should cease can be deduced. The threshold resource concentration for the expression of aggressive interference should concur with the intersection of the respective functional responses. Also, the intersection of Ni_IIIP_ and Ni_IIIλ_ corresponds with the threshold resource concentration, N_IIIcrit_:

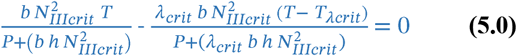

Solving equation **5.0** for N_III crit_, (assuming all parameters are positive) :

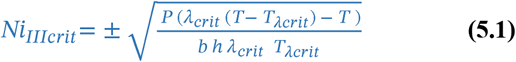

In equation **5.1**, the threshold time dedicated to aggressive interference (i.e., T_λcrit_) and threshold leverage (i.e., λ_crit_) are explicit . Solving equation **5.0** for these critical parameters yields the thresholds for time dedicated to aggressive interference (T_λcrit_ ) and leverage (λ_crit_):

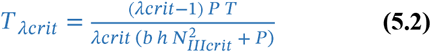

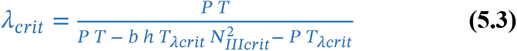

By definition, maximum time dedicated to aggressive behavior (i.e., T_λmax_) occurs at N_max_, so equation **3.0** also applies to the TypeIII functional response:

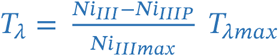

Upon rearrangement of equations **4.4** and **4.9**:

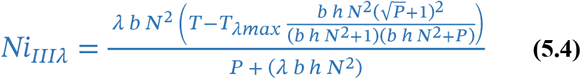

The threshold functional response value (Ni_IIIc rit_) corresponding to the threshold resource concentration (i.e., N_IIIcrit_) for the expression of aggressive interference for a Type III functional response occurs where there is no difference between equations **5.4** and **4.0**:

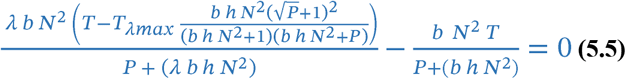

Solving expression **5.5** for Ni_IIIcrit_ :

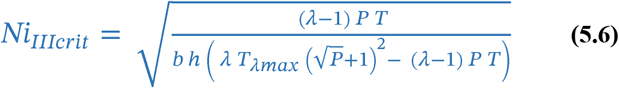

where Ni_IIIcrit_ stands for the threshold functional response value for which no difference exists between aggressive interference and equal access strategies. Equation **5.6** obviates resource concentration. Thus, Ni_IIIcrit_ denotes the point where aggressive interference and equal access functional responses intersect with respect to the threshold resource concentration, N_IIIt_. Implicitly, profitability for the equal access strategy exceeds that for the aggressive interference strategy at resource concentrations greater than N_IIIcrit_.

#### Real’s General Functional Response Model

A general functional response model based on enzyme kinetics equations for specifying reaction rates integrates the three standard functional response models along a continuum of model forms (Real 1977; 1979). Depending on the exponent, each of the standard models delineates a benchmark along the continuum. Instead of the squared exponent as in the Type III equation, the general model uses the global exponent (1 + q) to customize any functional response (i.e., q = 0 for Type II; q = 1 for Type III) (Rosenbaum and Rall 2018). Thus, the general functional response for a dominant consumer using aggressive interference as leverage can be cast in the form of the disc equation:

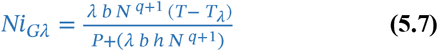

### Time spent on aggressive interference Implications

Expressing time spent on aggressive interference (T_λ_) in terms of other model parameters lends further insights into this tradeoff.

Given the following premises:

1. The time spent on aggressive interference or The time spent on dominant or subordinate aggressive interference behavior precludes time for obtaining resources (e.g., Ts, Th, etc.).
2. The difference between functional responses for individuals within *unrestricted* and *restricted* resource availability scenarios standardized by the maximum difference, 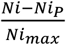, ranges as a proportion from 0 to 1:
  a. The difference = 0 at N = 0 and at N → ∞ (i.e., lim Δ = 0 at N → ∞ at Ni = T/h);
  b. The maximum difference between *Ni* and *Ni*_*P*_ defines Ni_max_;
  c. The peak value of 1 for the difference corresponds with the resource concentration, N_max_, for which time spent on aggressive interference is greatest ( *i. e., T*_λmax_).

Thus, the maximum proportion of time dedicated to aggressive interference occurring at resource concentration, N_max_:

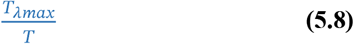

For any level of resource availability the product of three terms specifies time dedicated to aggression, T_λ_ . Because the form of the T_λ_ curve should reflect the normalized difference between per capita functional response s for *unrestricted* exclusive access and *restricted* equal access consumers :

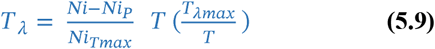

Cancelling the terms for total time in equation **5.9** yields the generalized form of the relationship inferred in equation **3.0** for specifying time spent on aggressive interference :

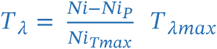

Accordingly, for any resource concentration (i.e., N), the time spent on aggressive interference scales T_λmax_ by the normalized difference between per capita functional response s for *unrestricted* exclusive access and *restricted* equal access consumers.

The time spent on aggressive interference must vary with certain key parameters that modulate the functional response of an aggressive consumer. Both consumer density (P) and leverage ( should help explain the time spent on aggressive interference :

- Higher P ≈ higher T_λmax_
- Higher λ ≈ lower T_λmax_

Thus, the proportion of time spent on aggressive interference (T_λmax_/T) can be exemplified by a function, *f*(*A*), incorporating effects of P and λ:

1. 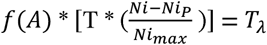, which varies dependently with N.
2. *f*(*A*) equates to the maximum proportion of time spent on aggressive interference, or 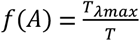.
3. As the product of two proportions, 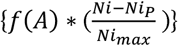 will also vary between 0 and 1:
  a. So, *f*(*A*) is a function that expresses the proportion of time spent on aggressive interference in terms of relevant functional response parameters i *f*(*A*) does not vary with resource availability, N; ii *f*(*A*) varies with λ and P;
  b. 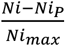 ranges between 0 and 1 with resou rce availability, N;

i. The forgoing expression for *T*_λ_ applies to any type of functional response:

1. assuming Ts, Th, and *T*_λ_ are exclusive.

### The leverage function (f(A))

The proposed function *f*(*A*) relates the counteracting effects of leverage (λ) and consumer density (i.e., P-1) on the functional response of a focal consumer exhibiting aggressive interference as leverage. Because λ reflects the efficacy of aggressive interference, it should vary inversely with time dedicated to such behavior. On the other hand, time dedicated to aggressive interference must increase with the density of contenders (i.e., P-1) within the ambit of the aggressive consumer.

The reciprocal exponential function fulfills the stipulation that the function, *f*(*A*), varies between the limits 0 and 1:

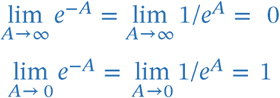

The parameters λ and P should affect *f*(*A*) in the following manner:

1. the leverage effect on *f*(*A*) ranges from 1 when λ is ineffectual to 0 when λ is completely effective (i.e., *T*_λ_ = T at λ = 0; *T*_λ_ = 0 at λ = ∞); and
2. the effect of P on *f*(*A*) ranges from 0 at (P-1) = 0 (i.e., solitary consumer) to 1 at (P-1) = ∞ (i.e., *T*_λ_ = 0 at (P-1) = 0; *T*_λ_ = T at (P-1) = ∞);
3. because *f*(*A*) = (1 − *e*^−*A*^), *f*(*A*) = 0 at *e*^−*A*^ = 1, and *f*(*A*) = 1 at *e*^−*A*^ = 0.

So:

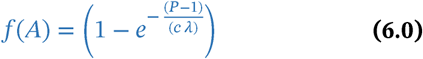

Where 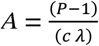. The scaling factor, c, transforms the joint effects of λ and P into a suitable form for yielding T_λmax_/T .

Furthermore, because:

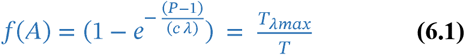

then:

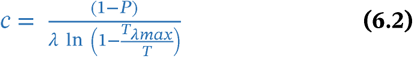

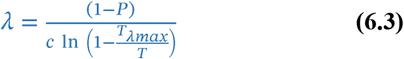

***Note***: Because both (1P) and ln (1- T_λmax_ /T) define negative values, both c and λ define positive values. And given both c and λ vary inversely with T_λmax_ (i.e., a lower T_λmax_ yields a higher (1- T_λmax_ /T) as well as a higher ( i.e., not as negative) ln (1- T_λmax_ /T) value ), the maximum proportion of time dedicated to aggressive interference decease s with both c and λ.

### Joint consumption within a group for a Type II functional response under sustained resource availability

Characterizing joint consumption by an aggressive agent within a group of consumers for a Type II functional response entails the synthesis of scenarios for the aggressive consumer and for the non-aggressive consumers. Firstly, t he encounter rate is partitioned proportionately within the group, so that it comprises terms representing the aggressive agent as well as the P-1 other consumers within a *restricted* scenario.

Thus,

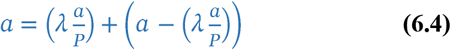

For an aggressive consumer within a group setting, the encounter rate:

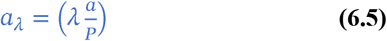

Rearrangement of the second term in **6.4** followed by normalization by P-1 yields the per capita encounter rate for a focal non-aggressive consumer:

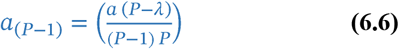

So, the compound difference equation for the joint Type II functional response under a sustained *restricted* resource availability scenario:

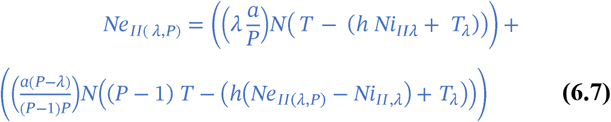

Where Ni_IIλ_ represents consumption by the aggressive consumer, T_λ_ denotes time dedicated to interference behavior, and Ne_II(λ,P)_ represents total consumption by the group, including the aggressive consumer. Furthermore, T_λ_ is spread equally among the P-1 non-aggressive consumers to account for the deficit in time available for consumption. The left hand term in equation **6.7** is identical to equation **1.8** for the aggressive agent; whereas differences between the right - hand term of equation **6.7** and equation **1.6** for the other consumers include effects of aggressive interference on encounter rates, reduced time available for consumption, and the specification of P-1 rather than P consumers (i.e., excluding the aggressive consumer).

Solving equation **6.7** for Ne_II(λ,P)_ yields the following equation:/

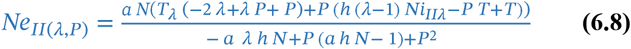

Conversely solving equation **6.7** for Ni_IIλ_ yields the following equation for obtaining Ni_IIλ_ :

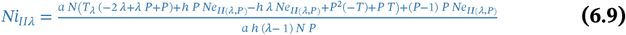

For any sustained level of resource availability (N) and set of model parameters, expected consumption by the focal aggressive consumer as well as by the non-aggressive consumers can be obtained and compared to actual values obtained as a test of the model. Estimation of expected values for unknown model parameters based on experiments can further inform equation **6.7**.

### Joint consumption within a group for a Type III functional response under sustained resource availability

Like for Type II functional response, characterizing joint consumption by an aggressive agent within a group of consumers for a Type III functional response entails the synthesis of scenarios for the aggressive consumer and for the non aggressive consumers. For a scenario involving sustained resource availability, the dynamic encounter rate, a=bN, is partitioned proportionately within the group. The dynamic encounter rate (bN) comprises terms for the aggressive agent as well as for the P-1 other consumers within a *restricted* scenario.

Thus,

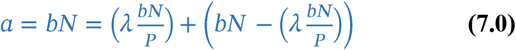

For an aggressive consumer within a group setting, the encounter rate:

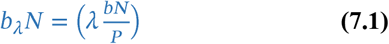

When parameters b, λ, N, and P are positive:

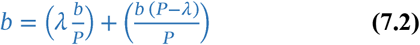

Rearrangement of the second term in **7.0** followed by normalization by P-1 yields the per capita encounter rate for a focal non-aggressive consumer:

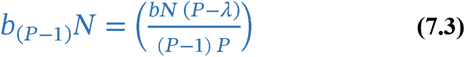

So, the compound difference equation for the joint Type III functional response under a sustained *restricted* resource availability scenario:

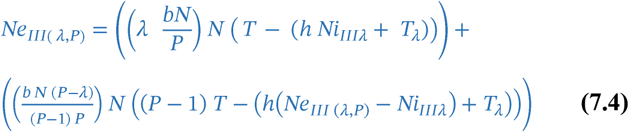

Where Ni_IIIλ_ represents consumption by the aggressive consumer, T_λ_ denotes time dedicated to interference behavior, and Ne_III(λ, P)_ represents total consumption by the group, including the aggressive consumer. Furthermore, T_λ_ is spread equally among the P-1 non-aggressive consumers to account for the deficit in time available for consumption. The left - hand term in equation **7.4** is identical to equation **4.3** for the aggressive agent ; whereas differences between the right - hand term of equation **7.4** and equation **3.9** for the other consumers include effects of aggressive interference on encounter rates, reduced time available for consumption, and the specification of P-1 rather than P consumers (i.e., excluding the aggressive consumer).

Solving equation **7.4** for Ne_III(λ,P)_ yields the following equation:

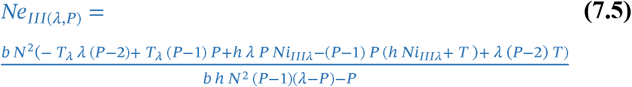

For any sustained level of resource availability (N) and set of model parameters, expected consumption by the focal aggressive consumer as well as by the non-aggressive consumers can be obtained and compared to actual values obtained as a test of the model. Estimation of expected values for unknown model parameters based on experiments can further inform equation **7.4**.

### Joint consumption within a group for a Type II functional response under depleted resource availability

Consumption by a group of agents often proceeds during diminishing resource availability, formally known as resource depletion (Rosenbaum and Rall 2018). While typically observed under experimental conditions, resource depletion also should frequently occur under natural conditions. Because the rate of consumption depends on resource availability, under resource depletion consumption decreases nonlinearly and continuously with time and resource concentration (Rogers 1972). Ordinary differential equation (ODE) methods can resolve the ensuing dynamic functional response under resource depletion (Rosenbaum and Rall 2018). Instead of a static expected rate of consumption expected under a sustained resource availability scenario, the resource depletion scenario integrates consumption over a specific period marked by known starting and ending resource concentrations.

An analytical solution to ODE known as the Rogers’ Random Predator Equation (RRPE) describes the Type II Functional Response under a depleted resource availability scenario (Royama 1971; Rogers 1972):

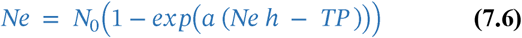

where Ne signifies the cumulative consumption realized at the end of an observation period, and N_0_ denotes the starting resource concentration. A practical method for explicitly solving the RRPE employs th e Lambert W function, or the product logarithm function (henceforth referred to as the product log function) (Rosenbaum and Rall 2018). The generalized product log function, *f(w) = we* ^*w*^, specifies the product of any complex number, *w*, and the exponential function, *e*^*w*^, raised to the power of the same complex number. In terms of z and w, *we*^*w*^ *= z* . The product log function has been used to describe various dynamic biological processes, including population growth (Corless et al. 1996), epidemiology (Lehtonen 2015), functional responses of predators and parasitoids (Arditi 1983), and microhabitat-related variability in the functional responses of crabs (Lipcius and Hines 1986). Similarly, the functional response involving an aggressive consumer within a group under resource depletion should be amenable to the product log function .

Equation **7.7** specifies the Type II functional response in RRPE form for P consumers assuming *unrestricted* encounter under a resource depletion scenario:

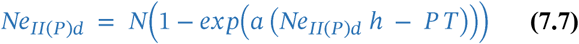

The following product log function estimates the solution for equation **7.7**:

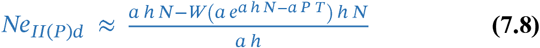

In contrast, equation **7.9** specifies the RRPE expressed as per capita consumption for a group of P consumers assuming *restricted* encounter under a resource depletion scenario:

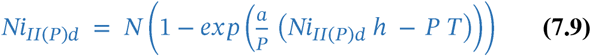

An alternative form of equation **7.9**:

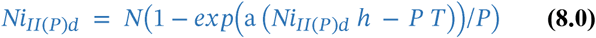

The estimated product log function solution for equations **7.9** and **8.0**:

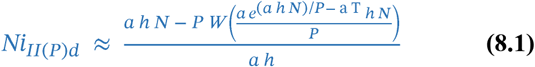

Equation **8.1** provides a reference for per capita consumption within a group of P consumers assuming *restricted* encounter under a resource depletion scenario.

Recall, the joint functional response assuming *restricted* encounter under a sustained resource availability scenario combines terms for the aggressive agent and for P-1 other consumers (i.e., equation **6.7**). Accordingly, equation **8.2** gives the RRPE equation for an aggressive agent and P-1 other consumers assuming *restricted* encounter under a resource depletion scenario:

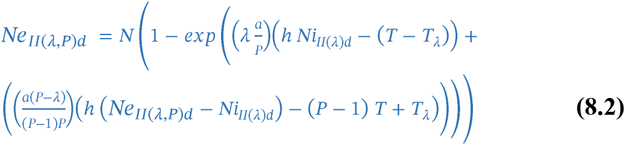

Given the initial resource concentration, N, and a set of parameter values, solutions of equation **8.2** yield expected consumption by the aggressive agent (Ni_II (λ)d_) as well as total consumption ( Ne_II(P)d_ ) (Fig. 6). Solving equation **8.2** given Ne_II(P)d_ provides a solution for the aggressive agent:

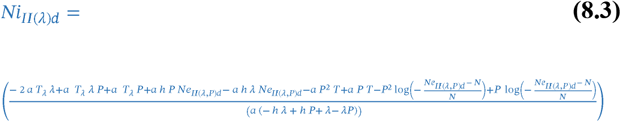

**Figure 6.**
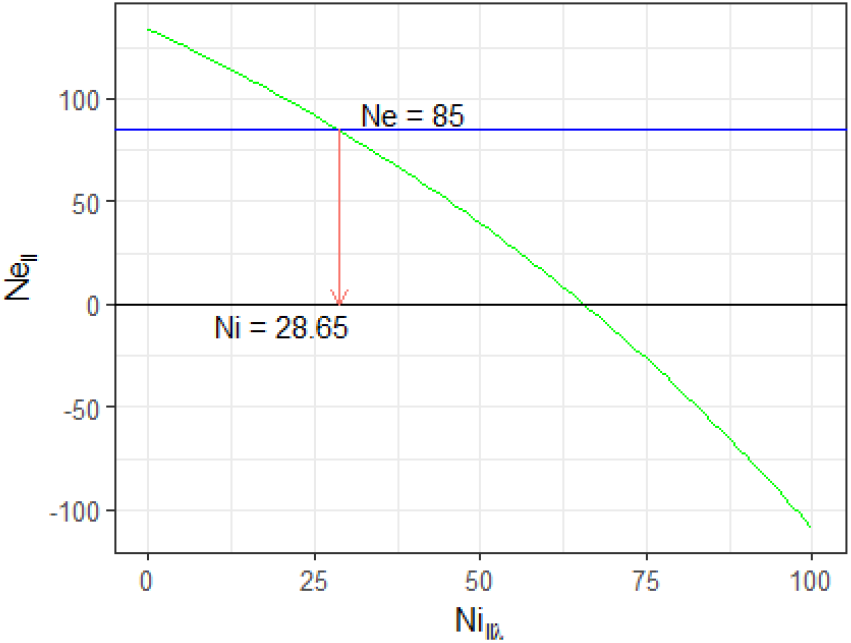
Joint consumption within a group for a Type II functional response under depleted resource availability. Graphic solution of the product log function for expected consumption while resource availability is progressively depleted. Here, total consumption by the group (Ne_II(P)_) is 85 and consumption by the aggressive agent (Ni_II(λ)_) is 28.65. For this depletion curve, a = 0.2, h = 0.09, P = 5, T = 4, λ = 3, and T_λ_,= 0.4. Per capita consumption by the P-1 other consumers, Ni_II(P)_ = (Ne_II(P)_ Ni_II(λ)_)/P-1, or (85-28.65)/4 = 14.09. (a = resource encounter rate; h = handling time per unit resource; P = number of consumers; T = total time; λ = leverage; T_λ_ = time spent on aggression).

Solving equation **8.2** given Ni_II (λ)d_ also yields the product log equation specifying total consumption by the aggressive agent and the P-1 other consumers under resource depletion:

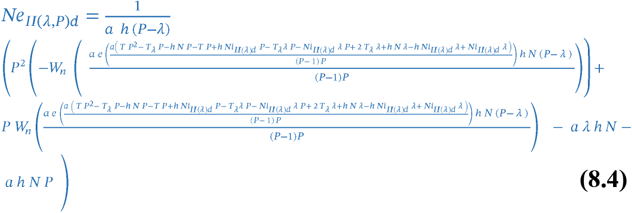

Consumption by the other P-1 consumers can then be obtained by subtraction (i.e., Ne_II( λ, P)d_ - Ni_II( λ)d_), followed by division by P-1 to obtain the per capita cumulative consumption for non-aggressive consumers at time T under resource depletion. Equation **8.2** can also provide solutions for unknown parameters, such as the leverage coefficient, λ.

### Joint consumption within a group for a Type III functional response under depleted resource availability

Because the Type III functional response entails a dynamic resource-dependent encounter rate, Rosenbaum and Rall (2018) presented a corrected product log function for estimating the Type III functional response under resource depletion, based on a simplified form of the version by Okuyama and Ruyle (2011). Consequently, the RRPE equation yielding the correct log product function for the Type III functional response under resource depletion is analogous to the RRPE equation for the Type II functional response:

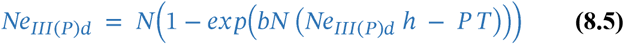

The following product log function estimates the solution for equation **8.5**:

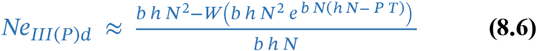

Equation **8.6** compares favorably with the product log function given in Rosenbaum and Rall (2018) for the Type III functional response as expressed for P consumers.

In contrast, equation **8.7** specifies the RRPE expressed as per capita consumption for a group of P consumers assuming *restricted* encounter under a resource depletion scenario:

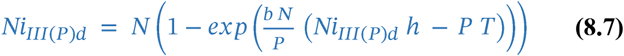

An alternative form of equation **8.6**:

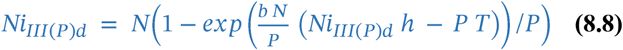

The estimated product log function solution for equations **8.7** and **8.8**:

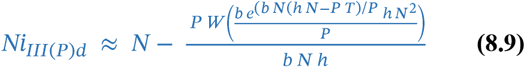

Equation **8.9** provides a reference for per capita consumption within a group of P consumers assuming *restricted* encounter under a resource depletion scenario.

Recall, the joint functional response assuming *restricted* encounter under a sustained resource availability scenario combines terms for the aggressive agent and for P-1 other consumers (i.e., equation **7.4**). Accordingly, equation **9.0** gives the RRPE equation for an aggressive agent and P-1 other consumers assuming *restricted* encounter under a resource depletion scenario:

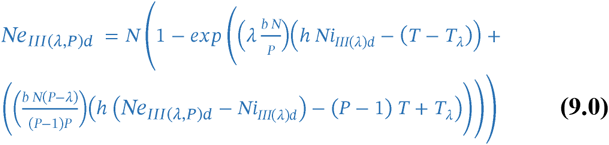

Given the initial resource concentration, N, and a set of parameter values, solutions of equation **9.0** yield expected consumption by the aggressive agent (Ni_III (λ)d_) as well as total consumption ( Ne_III(P)d_ ). Solving e quation **9.0** given Ne_III(P)d_ provides a solution for the aggressive agent:

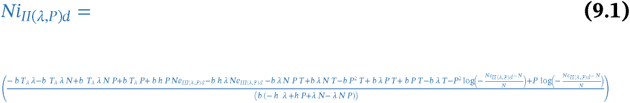

When given Ni_III(λ)d_, equation **9.0** also yields the product log equation specifying the total cumulative consumption under resource depletion. Consumption by the other P-1 consumers can be obtained by subtraction (i.e., Ne_III( λ, P)d_ Ni_III( λ)d_), followed by division by P-1 to obtain the per capita cumulative consumption for non-aggressive consumers at time T under resource depletion. Finally, equation **9.0** can provide solutions for unknown parameters, such as the leverage coefficient, λ.

## Discussion

The proposed functional response models envisage how aggression should be effectively used as leverage to boost the rate of resource consumption. The models project the intensity of aggression and the functional responses of consumers under varying levels of resource availability and consumer density, given leverage effects of aggression. As such, the models explicitly comprise the costs and benefits of interference in terms of time spent on aggression relative to resources gained. Herein, aggressive interference manifests as an asymmetric mechanism for increasing the rate of consumption above the level specified by a premise of equal access, as a standard of comparison. Consequently, leverage elevates the realized rate of gain for aggressive consumers while suppressing the rate of gain for non-aggressive consumers within a group. In addition, the proposed models specify differential expected consumption for aggressive and non-aggressive consumers within groups for both sustained and depleted resource scenarios. Thus, the proposed models provide a framework for predicting collective functional responses for various configurations of aggressive and non-aggressive consumers.

Consumer-dependent functional response models conventionally treat interference as a mass-action effect (Hassell 1978; Van Der Meer and Smallegange 2009; Novak and Stouffer 2021), wherein the collective functional response is lowered through mutual distraction (Novak and Stouffer 2021). Some functional response studies also recognize and document the importance of unequal, or asymmetric interference (Abrams and Ginzburg 2000; Nillson et al. 2004). By modifying conventional interference coefficients, Nillson et al. 2004 distinguished unequal interference behavior among juvenile salmonids for which collective functional responses of dominant or subordinate fishes were elevated or depressed, respectively. Moreover, functional responses for solitary consumers reached higher levels than those for either dominant or subordinate consumers together. Validation of interference functional response models generally entails empirical regression methods, where better fits with interference imply increased realism (Skalski and Gilliam 2001). However, existing functional response models lack explicit mechanistic explanations of interference (Nillson et al. 2007). This work addresses this deficit by explicitly incorporating relevant parameters and assumptions into functional response models at the individual level from which more inclusive levels can be envisaged.

Other sorts of models consider asymmetric effects of aggression for elevating resource gain using different analytical methods, including game theory (Giraldeau and Caraco 2000; Sirot 2000; Dubois and Giraldeau 2005), and individual-based models (Stillman et al. 1997). The latter uses optimal decision theory to mediate asymmetric behavioral interactions, such as kleptoparasitism. Previous studies confirm that consumers express aggressive interference facultatively, depending on changing ecological trade-offs (Stillman et al. 1997; Goss-Custard et al. 1998, as cited in Sirot 2000). The Dove-Hawk game theory model also confirms the evolutionary stability of a flexible aggressive interference strategy (sensu Smith and Price 1973; Cowden 2012; Křivan and Cressman 2017. Indeed, time spent during an interaction modulates whether a consumer assumes the hawk strategy, despite a net benefit for employing that strategy (Křivan and Cressman 2017). Previous studies also confirm other assumptions and predictions made in this study. Resource-dependent interference should wane as the functional response saturates with increasing resource availability (Skalski and Gilliam 2001). Furthermore, aggression intensity has been shown to vary non-linearly with respect to group size (Dubois and Giraldeau 2005). Importantly, a threshold consumer density for which the net benefit of aggression equals zero has been demonstrated (Stillman et al. 1996). Mechanistic consumer-dependent functional response models should help clarify forgoing phenomena within a coherent framework.

The proposed functional response models yield several falsifiable predictions, including the resource concentration at which aggressive interference should be greatest, as well as the functional response value and corresponding threshold resource concentration above which aggressive interference should cease to be effective. The intensity of aggression peaks at some intermediate level of resource availability within the proposed models in connection with the scope for gain conveyed by the normalized difference between per capita functional response s for *unrestricted* exclusive access and *restricted* equal access consumers . In contrast, the intensity of aggressive behavior varies inversely with resource availability within the Hawk-Dove game theory model of aggressive interference, which predicts the probabilistic outcomes of multiple one-to-one conflicts based on costs in time and energy (Sirot 2000). This prediction has been used in support of aggression as a reflection of competition in the Apennine Chamois when resources become limited (Fattorini et al. 2018). The Dove-Hawk model focuses on the evolutionary stability of the strategy whereas the proposed functional response models ensue during an ecological time frame. Nevertheless, this clear distinction in the expression of aggression with respect to resource availability provides a means to discern which model applies for a given case.

When unconstrained, the per capita rate of consumption reaches the level afforded by sustained resource availability. However, other consumers constrain the rate of consumption an aggressive consumer can reach within the proposed functional response models. The volume /area potentially under control of the aggressive consumer for garnering resources defines its ambit, or ‘area of discovery.’ And leverage can effectively deter multiple contenders concurrently within the ambit, while also precluding the aggressor from directly obtaining resources. For a fixed ambit, space may become the *de facto* resource when control by the aggressor extends to territorial defense. However, the ambit would typically move with a mobile aggressive consumer as it seeks resources. Although the actual contenders within the ambit might vary from moment to moment, the effective number of overlapping consumers should stabilize over a long enough period. Furthermore, an implicit trade-off exists in which some consumers may access the ambit while the aggressor actively suppresses others. Such implicit features of the proposed functional response models stand in contrast to game-theory models involving consecutive one-to-one contests (Sirot 2000). Which type of model applies likely depends upon the nature of behavioral interactions and the time frame under consideration.

The extent to which leverage deters contenders depends on how strongly aggressive interference is perceived as a threat. Aggressive behavior often appears in conjunction with associated traits that evolved for intimidation (King 1973). The “ecology of fear” subfield emerged from studies of how the threat of predation indirectly suppresses prey behavior through multifarious cues (Brown et al. 1999). For example, trophic cascades occur when a top predator indirectly suppresses feeding by an intermediate predator on a basal prey (Peacor et al. 2020; Schweiss and Rakocinski 2020). The threat of predation also induces the development of social groups of foragers ( Giraldeau and Caraco 2000). Aggressive members of such groups should benefit at the expense of subordinate members. Aggressive interference might alleviate predation intensity by distracting consumers from feeding, as has been noted in studies of mutual interference (Hassell 1978). Even where social foraging enhances prey encounter rates ( Giraldeau 1984), a dominant consumer might reach even higher rates of consumption within the group due to benefits of learning or shared vigilance.

Several assumptions may limit the applicability of the proposed aggressive interference functional response models. In the proposed models, time is spent on dispensing or evading aggression exclusively of time spent on consumption. Also, the proposed models assume fixed leverage with respect to consumer density and resource availability. In standard models, fixed encounter rate or handling time parameters sometimes prove unrealistic (Abrams 1990; Abrams 2010; Gibert and DeLong 2015), and instead may vary with consumer density or resource concentrations (Okuyama 2010; Okuyama 2012), or with changing environmental variables or resource traits ( Robertson and Hammill 2021). Indeed, the standard Type III functional response model incorporates a dynamic encounter rate to account for improved consumption skills with experience in connection with resource availability (Holling 1966). Similarly, handling time may decrease with respect to resource availability (Abrams 1990) or vary among individual consumers (Hartley et al. 2019). Dynamic parameters can be used in the interest of realism where the proposed simple models reveal the faulty use of static parameters (*sensu* Okuyama 2012 ). For example, leverage might vary systematically with resource or contender density . Experience might reduce time dedicated to aggressive interference due to improved leverage efficacy or enhanced avoidance by subordinate consumers (Stillman et al. 1997).

Predictions following from the proposed aggressive interference functional response models can be tested by replicated experiments involving quantification of consumption and aggression by a dominant focal subject together with several subordinate consumers within the same size class. Both general and specific predictions following from the models could be tested. For example, a previous experiment employing a dominance hierarchy of bluegill sunfish (*Lepomis macrochirus* ) supported the general expectation that the frequency of aggressive interference followed a hump shaped nonlinear relationship with respect to depleting food availability (Rakocinski et al. 1983). Moody and Ruxton (1996) also confirmed a hump-shaped non-linear relationship in the frequency of aggression with respect to resource availability. Recorded observations of per capita consumption using video would facilitate tests of more specific predictions. Uncertainty estimates for paramete rs require replicated observations made at various sustained levels of resource availability. Key variables include rates of consumption for the focal aggressive as well as subordinate consumers. In addition, quantification of handling time (T_h_), encounter rates (a), and time spent on aggression (T_λ_), would enable estimation of time spent while obtaining and searching for resources.

Critical features of the proposed models are amenable to empirical testing through controlled experiments. Expected aggression intensity with respect to resource concentration could be obtained from the difference between the *ad-libitum* functional response of the solitary aggressive consumer and the per capita functional response for subordinate consumers. The resource concentration at which peak aggression (T_λmax_) should occur would follow. Additionally, whether aggression ceases at the predicted level of resource availability where the functional response of the aggressive consumer equals that of the per capita subordinate consumer could be discerned. Lambda ( λ) could be estimated from comparisons of the realized functional responses of the focal aggressive consumer and subordinate consumers. Extraneous effects of satiation or other variables on the motivation of subjects might also undermine the assumptions of the proposed models. Experiments could also test whether cumulative consumption by focal aggressive and subordinate consumers agrees with expectations based on the proposed depletion models. Ultimately, one may determine whether the data are better explained by alternative hypotheses involving mutual interference as opposed to aggressive interference.

Although models can help explain phenomena by ruling out alternative explanations (Ylikoski and Aydinonat 2014), simplified mathematical relationships falsely idealize complex natural phenomena. For example, while the proposed models characterize the cumulative time dedicated to aggression, they do not specify the dispersion of aggressive and consumptive behaviors integrated over the period spent consuming resources. Presumably, the aggressor would perform tactically with respect to the dispersion of resources and consumers on shorter time scales. Also, the efficacy of aggressive interference behavior for garnering resources depends on how appropriately coupled stimuli act to induce the behavioral responses (Bindra 1978, Zaman et al. 2023). Natural selection should favor the accurate expression of aggressive behavior to enhance resource gain due to the direct connection with fitness (Georgiev et al. 2013). Natural selection should also minimize risks that could reinforce the efficacy of aggressive interference. For example, consumers at lower positions within a dominance hierarchy face greater risk of mortality from compounded effects of aggressive interference. They also face combined effects of slower growth, greater injury, and higher predation. Such auxiliary effects would reinforce selection for effectively wielding leverage through aggressive interference to maximize resource gain. Resultant den sity-dependent effects of aggressive interference should also mediate spacing of consumers, reproductive outputs, and population structure.

The proposed models of aggressive interference define functional responses for which a focal aggressive consumer elevates its intake of resources via equilaterally distributed leverage among any number of subordinate consumers with equal access to resources. The model scenarios presented heuristically exemplify simplified straightforward dominance-subordinate configurations. However, fully developed dominance hierarchies form cascading networks of behavioral interactions (Cumming 2016; Fattorini et al. 2018 ). Thus, groups often embody complex hierarchies wherein the same individuals may display both aggression and evasion. Such complex hierarchical configurations can be formulated within a framework defined by the same precepts as those used in this study. For example, the overarching trait of body size frequently confers position-related polarity within a dominance hierarchy; thus, leverage within a group may also covary in conjunction with body size ( Mesterton-Gibbons and Dugatkin . 1995). But aggressive interference might also decrease with disparity in body size (Woodward et al. 2005), due to corresponding non-overlap in resource requirements. Furthermore, better foraging efficiency may promote higher positions within dominance hierarchies, notwithstanding body-size disparities (Hartley et al. 2019).

More realistic and complex aggressive interference models using the same precepts as those in this paper could prove computationally challenging. A general systematic framework for specifying diverse pairwise interactions could employ matrix calculus along with optimization methods such as linear and nonlinear programming. Such an approach would also be conducive to extended models based on relaxed parameter assumptions. Modified models could incorporate flexible leverage parameters among consumers within a dominance-hierarchy. Matrix-based methods could also aid in the formulation of stochastic aggressive interference functional response models by accommodating variance structures for parameters, etc. Stochastic forms of deterministic functional response models have been developed (Gilioli et al. 2008; Van Der Meer and Smallegange 2009; Palamara et al. 2021). An exemplary study has used maximum likelihood estimation of parameters of the Beddington functional response model of mutual interference among a small number of consumers (Van Der Meer and Smallegange 2009). Finally, in addition to consumption, resource availability and aggressive interference could be modeled stochastically.

## Conclusions

Assuming aggressive behavior acts as a leverage mechanism for boosting resource gain above what would be obtained under a premise of equal access, the expression of asymmetric interference by a focal aggressive consumer should correspond with the potential for resource gain as determined by tradeoffs involving leverage strength, resource availability and consumer density. Proposed models expand on these relationships to make falsifiable predictions pertaining to the expression of aggressive interference with respect to various tradeoffs:

1. Assumng the difference between the functional response for a solitary consumer with exclusive access and that for a consumer within a group with equal access to a *restricted* resource drives the intensity of aggressive interference, the maximum intensity of aggressive behavior should occur at the resource concentration equivalent to 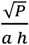 for a Type II, or 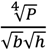 for a Type III functional response.
2. The resource concentrations and functional response values above which aggressive interference will not effectively act as leverage occurs at explicitly defined resource concentration and functional response thresholds for Type II and Type III functional responses. The functional response value below which aggressive interference should not gainfully provide leverage occurs at explicitly defined thresholds for Type II and Type III functional responses.
3. A mechanistic interpretation of the maximum proportion of time dedicated to aggressive interference (i.e., 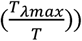 can be expressed in terms of leverage and consumer density .
4. When encounter rates are correctly normalized, an equation for joint consumption by an aggressive agent can be additively combined with terms for any number of other consumers under sustained *restricted* resource availability for Type II and Type II functional responses.
5. When encounter rates are correctly normalized, the Rogers’ Random Predator Equation (RRPE ) for joint consumption by an aggressive agent can be additively combined with terms for any number of other consumers under depleting resource availability for Type II and Type II functional responses. The RRPE equations yield product log solutions for expected consumption by the aggressive agent as well as for the other consumers under resource depletion.

## Acknowledgements

Thanks go to Wolfram Research Inc., Champaign, IL for open access to the Wolfram|Alpha knowledge engine which routinely provided computational ground truthing support. This work was conceived while employed as a faculty member of the University of Southern Mississippi.

